# Myomatrix arrays for high-definition muscle recording

**DOI:** 10.1101/2023.02.21.529200

**Authors:** Bryce Chung, Muneeb Zia, Kyle A. Thomas, Jonathan A. Michaels, Amanda Jacob, Andrea Pack, Matthew J. Williams, Kailash Nagapudi, Lay Heng Teng, Eduardo Arrambide, Logan Ouellette, Nicole Oey, Rhuna Gibbs, Philip Anschutz, Jiaao Lu, Yu Wu, Mehrdad Kashefi, Tomomichi Oya, Rhonda Kersten, Alice C. Mosberger, Sean O’Connell, Runming Wang, Hugo Marques, Ana Rita Mendes, Constanze Lenschow, Gayathri Kondakath, Jeong Jun Kim, William Olson, Kiara N. Quinn, Pierce Perkins, Graziana Gatto, Ayesha Thanawalla, Susan Coltman, Taegyo Kim, Trevor Smith, Ben Binder-Markey, Martin Zaback, Christopher K. Thompson, Simon Giszter, Abigail Person, Martyn Goulding, Eiman Azim, Nitish Thakor, Daniel O’Connor, Barry Trimmer, Susana Q. Lima, Megan R. Carey, Chethan Pandarinath, Rui M. Costa, J. Andrew Pruszynski, Muhannad Bakir, Samuel J. Sober

## Abstract

Neurons coordinate their activity to produce an astonishing variety of motor behaviors. Our present understanding of motor control has grown rapidly thanks to new methods for recording and analyzing populations of many individual neurons over time. In contrast, current methods for recording the nervous system’s actual motor output – the activation of muscle fibers by motor neurons – typically cannot detect the individual electrical events produced by muscle fibers during natural behaviors and scale poorly across species and muscle groups. Here we present a novel class of electrode devices (“Myomatrix arrays”) that record muscle activity at unprecedented resolution across muscles and behaviors. High-density, flexible electrode arrays allow for stable recordings from the muscle fibers activated by a single motor neuron, called a “motor unit”, during natural behaviors in many species, including mice, rats, primates, songbirds, frogs, and insects. This technology therefore allows the nervous system’s motor output to be monitored in unprecedented detail during complex behaviors across species and muscle morphologies. We anticipate that this technology will allow rapid advances in understanding the neural control of behavior and in identifying pathologies of the motor system.

## Introduction

Recent decades have seen tremendous advances in our understanding of the physiological mechanisms by which the brain controls complex motor behaviors. Critical to these advances have been tools to record neural activity at scale^4,5^, which, when combined with novel algorithms for behavioral tracking ^6–10^, can reveal how neural activity shapes behavior ^11,12^. In contrast, current methods for observing the nervous system’s motor output lag far behind neural recording technologies. The nervous system’s control of skeletal motor output is ultimately mediated by “motor units”, each of which consists of a single motor neuron and the muscle fibers it activates, producing motor unit action potentials (**Fig. 1a**) that generate muscle force to produce movement ^13^. Because each action potential in a motor neuron reliably evokes a single spike in its target muscle fibers, action potentials recorded from muscle provide a high-resolution readout of motor neuron activity in the spinal cord and brainstem. However, our understanding of motor unit activity during natural behaviors is rudimentary due to the difficulty of recording spike trains from motor unit populations.

Traditional methods for recording muscle fiber activity via electromyography (EMG) include fine wires inserted into muscles and electrode arrays placed on the surface of the skin ^14^. These methods can resolve the activity of individual motor units in only a limited range of settings. First, to prevent measurement artifacts, traditional EMG methods require that a subject’s movements be highly restricted, typically in “isometric” force tasks where subjects contract their muscles without moving their bodies ^15–18^. Moreover, fine wire electrodes typically cannot detect single motor unit activity in small muscles, including the muscles of widely used model systems such as mice or songbirds ^19–21^, and surface electrode arrays are poorly tolerated by freely behaving animal subjects. These limitations have impeded our understanding of fundamental questions in motor control, including how the nervous system coordinates populations of motor units to produce skilled movements, how this coordination degrades in pathological states, and how motor unit activity is remapped when animals learn new tasks or adapt to changes in the environment.

**Figure 1:**
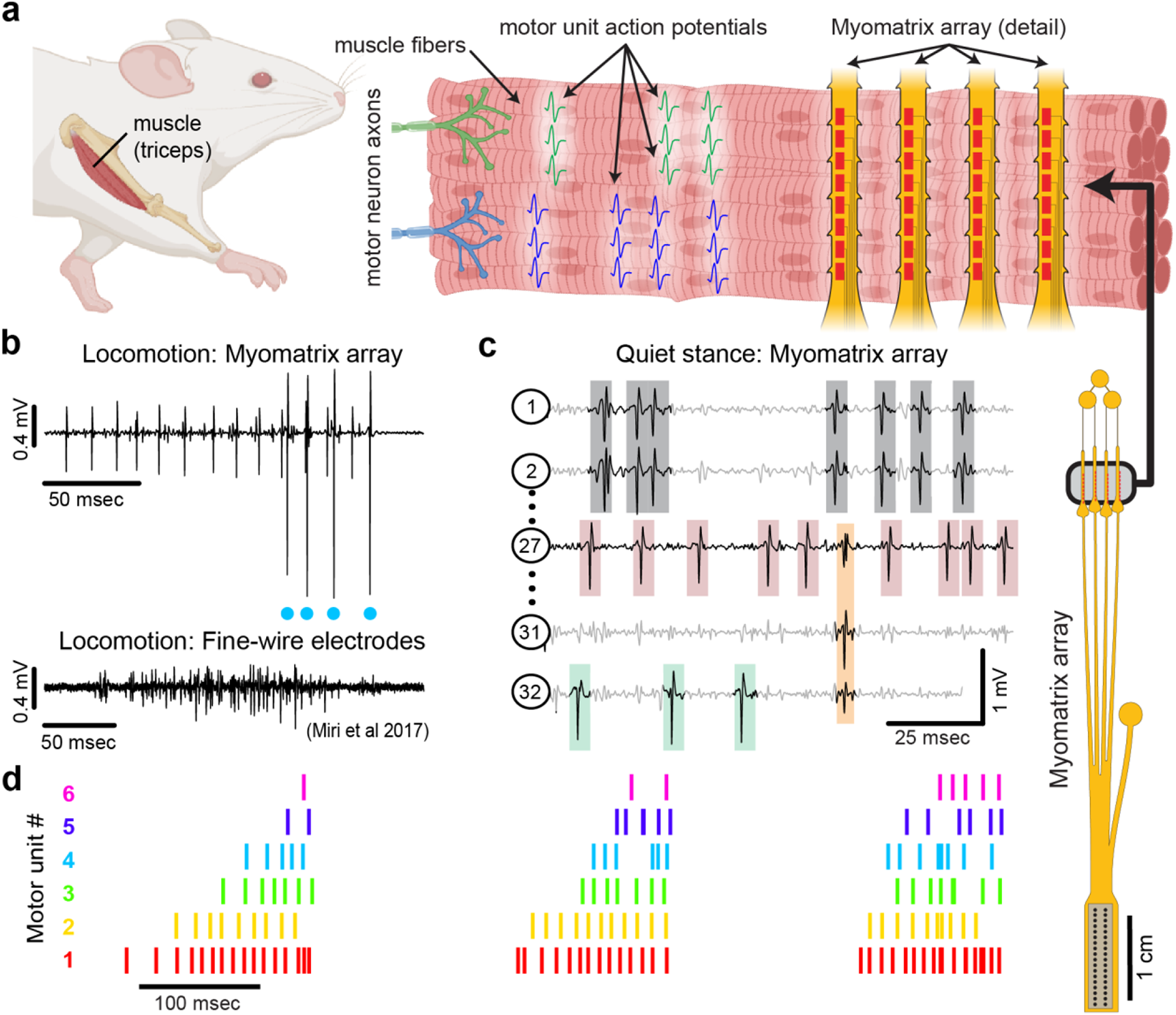
Myomatrix arrays record muscle activity at motor unit resolution. **(a)** The nervous system controls behavior via motor units, each consisting of a single motor neuron and the muscle fibers it innervates. Each motor neuron’s spiking evokes motor unit action potentials in the corresponding muscle fibers. Myomatrix arrays (right) bearing 32 electrode contacts on a flexible substrate (**Supplemental Fig. 1**) can be targeted to one or more muscles and yield high-resolution recordings of motor activity during free behavior. Motor neurons, muscle fibers, and electrode arrays are not shown to scale. **(b,c)** Example recordings from the right triceps muscle of a freely behaving mouse. **(b)** Top, bipolar Myomatrix recording from the mouse triceps during locomotion. Blue dots indicate the spike times of one motor unit isolated from the data using a spike sorting method based on principal components analysis (**Supplemental Fig. 2a-d**). Bottom, example data (from ^1^, used with permission) from traditional fine-wire EMG recording of triceps activity during locomotion. Applying the PCA-based spike sorting method to the fine-wire data did not isolate any individual motor units. **(c)** Unipolar Myomatrix recording during quiet stance. Colored boxes illustrate motor unit action potentials from four identified units. Spike waveforms from some units, including those highlighted with gray and orange boxes, appear on multiple electrode channels, requiring the use of a multi-channel spike sorting algorithm (Kilosort 2.5, see **Supplemental Fig. 2e-h**). **(d)** Spiking pattern (tick marks) of six individual motor units recorded simultaneously during locomotion on a treadmill. The three bursts of motor unit action potentials correspond to triceps activity during three stride cycles. Motor unit 4 (cyan) is the same motor unit represented by cyan dots in (b). The other motor units in this recording, including the smaller amplitude units at top in (b), were isolated using Kilosort but could not be isolated with the PCA-based method applied to data from only the single recording channel shown (b).

Here, we present a novel approach (**Fig. 1**) to recording populations of individual motor units from many different muscle groups and species during natural behaviors. Flexible multielectrode (“Myomatrix”) arrays were developed to achieve the following goals:

a. Record muscle activity at motor unit resolution
b. Record motor units during active movements
c. Record from a wide range of muscle groups, species, and behaviors
d. Record stably over time and with minimal movement artifact

To achieve these goals, we developed a variety of array configurations for use across species and muscle groups. Voltage waveforms from individual motor units (**Fig. 1b,c**) can be readily extracted from the resulting data using a range of spike-sorting algorithms, including methods developed to identify the waveforms of individual neurons in high-density arrays ^2,22^. Below, we show how Myomatrix arrays provide high-resolution measures of motor unit activation in a variety of species and muscle groups including forelimb, hindlimb, orofacial, pelvic, vocal, and respiratory muscles.

## Results

We developed methods to fabricate flexible, high-density EMG (“Myomatrix”) arrays, as detailed in the Methods and schematized in **Supplemental Figure 1**. We selected polyimide as a substrate material due to its strength and flexibility and the ease with which we could define electrode contacts, suture holes, and other sub-millimeter features that facilitate ease of implantation and recording stability (**Supplemental Fig. 1a-e**). Moreover, simple modifications to the fabrication pipeline allowed us to rapidly design, test, and refine different array morphologies targeting a range of muscle sizes, shapes, and anatomical locations (**Supplemental Fig. 1c, f, g**).

### Myomatrix arrays record muscle activity at motor unit resolution

Myomatrix arrays robustly record the activity of individual motor units in freely behaving mice. Arrays specialized for mouse forelimb muscles include four thin “threads” (8 electrodes per thread, 32 electrodes total) equipped with suture holes, flexible “barbs,” and other features to secure the device within or onto a muscle (**Fig. 1a, Supplemental Fig. 1c, d, e, h**). These devices yielded well-isolated waveforms from individual motor units (**Fig. 1b**, top), which were identified using open-source spike sorting tools ^2,22^. As detailed in **Supplemental Figure 2a-d**, in some cases the spike times of individual motor units (cyan dots, **Fig. 1b**) can be isolated from an individual electrode channel with simple spike sorting approaches including single-channel waveform clustering ^22^. In other cases, waveforms from individual motor units appeared on multiple electrode channels (**Fig. 1c**), allowing – and in many cases necessitating – more advanced spike-sorting approaches that leverage information from multiple channels to identify larger numbers of units and resolve overlapping spike waveforms ^2^, as detailed in **Supplemental Figure 2e-h**. These methods allow the user to record simultaneously from ensembles of single motor units (**Fig. 1c,d**) in freely behaving animals, even from small muscles including the lateral head of the triceps muscle in mice (approximately 9 mm in length with a mass of 0.02 g ^23^). Myomatrix recordings isolated single motor units for extended periods (greater than two months, **Supplemental Fig. 3e**), although highest unit yield was typically observed in the first 1-2 weeks after chronic implantation. Because recording sessions from individual animals were often separated by several days during which animals were disconnected from data collection equipment, we are unable to assess based on the present data whether the same motor units can be recorded over multiple days.

### Myomatrix arrays record motor units during active movements

Myomatrix arrays outperform traditional fine-wire electrodes in mice by reliably recording isolated single units in behaving animals. First, Myomatrix arrays isolate the activity of multiple individual motor units during freely moving behavior (**Fig. 1c-d**). In contrast, wire electrodes typically cannot resolve individual motor units during muscle lengthening and shorting, as occurs in naturalistic movements such as locomotion ^1,24^. **Figure 1b** illustrates a comparison between Myomatrix (top) and fine-wire (bottom) data recorded during locomotion in the mouse triceps. Spike-sorting identified well-isolated motor unit spikes in the Myomatrix data (cyan dots in **Fig. 1b, top**) but failed to extract any isolated motor units in the fine wire data (**Supplemental Fig. 2a,b**). Similarly, while Myomatrix recordings robustly isolated motor units from a songbird vocal muscle, fine wire EMG electrodes applied to the same muscle did not yield isolatable units (**Supplemental Fig. 2c,d**). This lack of resolution, which is typical of fine wire EMG, severely limits access to motor unit activity during active behavior, although wire electrodes injected through the skin can provide excellent motor unit isolation during quiet stance in mice ^25^. Second, because wire-based EMG requires inserting an additional wire for each additional electrode contact, only a single pair of wires (providing a single bipolar recording channel, **Fig. 1b**, bottom) can be inserted into an individual mouse muscle in most cases ^1,24,26^. In contrast, at least four Myomatrix “threads” (**Fig 1a**), bearing a total of 32 electrodes, can be inserted into one muscle (**Fig. 1c** shows five of 32 channels recorded simultaneously from mouse triceps), greatly expanding the number of recording channels within a single muscle. Single motor units were routinely isolated during mouse locomotion in our Myomatrix recordings (**Fig. 1**), but never in the fine-wire datasets from ^1^ we re-analyzed or, to our knowledge, in any prior study. Moreover, in multiunit recordings, Myomatrix arrays have significantly higher signal-to-noise ratios than fine-wire EMG arrays (**Supplemental Fig. 3**). Myomatrix arrays therefore far exceed the performance of wire electrodes in mice in terms of both the quality of recordings and the number of channels that can be recorded simultaneously from one muscle.

**Figure 2:**
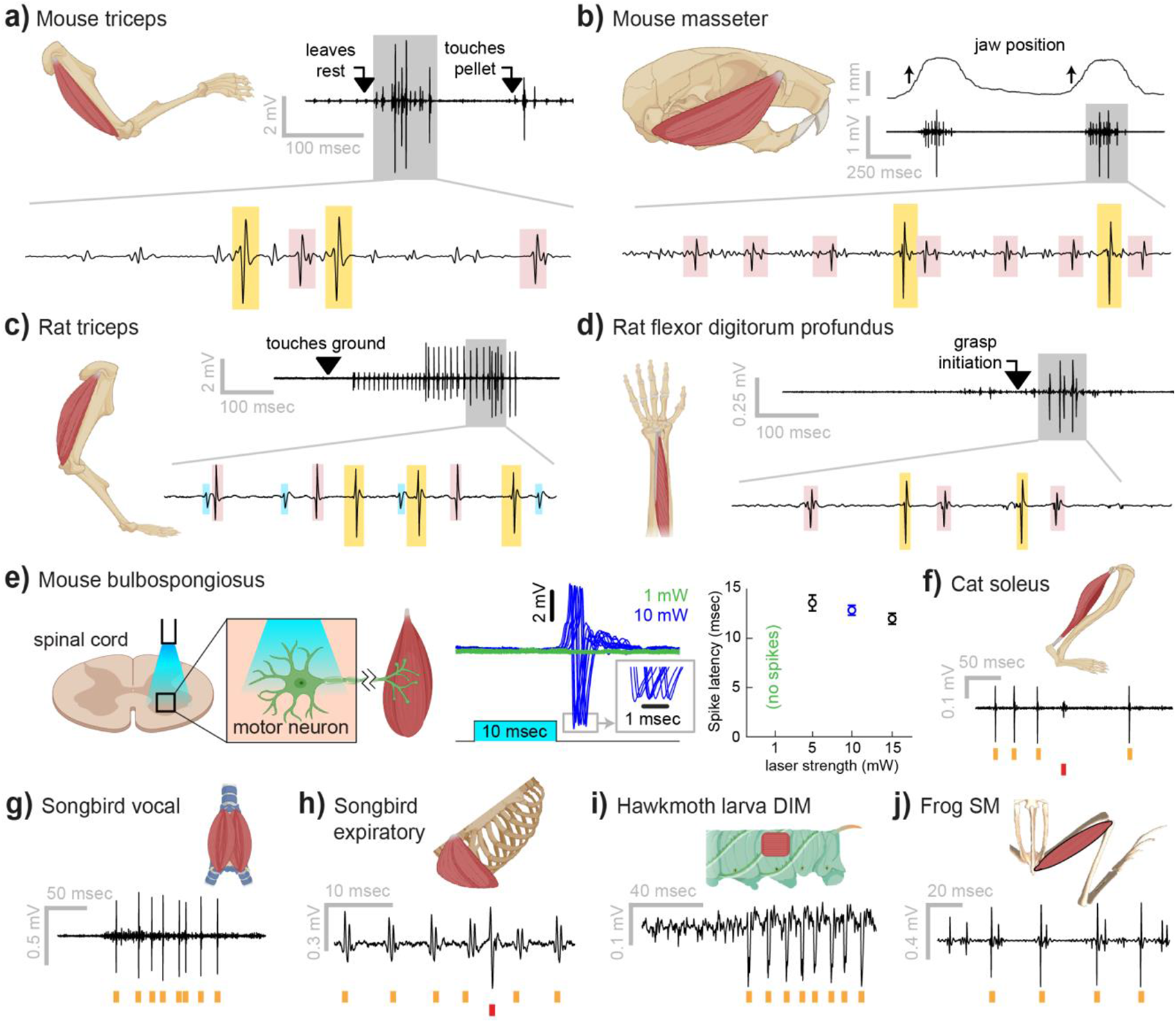
Myomatrix recordings across muscles and species. **(a)** Example recording from **mouse triceps** during a head-fixed pellet reaching task. Arrows at top indicate the approximate time that the animal’s paw leaves a rest position and first contacts the target. Bottom, colored boxes highlight motor unit action potentials identified using Kilosort ^2^. Different box colors on the same voltage trace indicate distinct motor units. **(b)** Recordings from the **mouse superficial masseter** muscle were obtained in anesthetized, head-fixed mice when passive mandible displacement evoked reflexive muscle contractions. Top trace shows the lateral component of jaw displacement, with arrows indicating the direction and approximate time of displacement onset. **(c)** In a recording from **rat triceps** during head-free locomotion, the arrowhead indicates the time that the mouse’s paw touched the treadmill surface, marking the beginning of the stance phase. **(d)** Recording from the **rat flexor digitorum profundus** muscle during a pellet reaching task, arrow indicates the time of grasp initiation. **(e)** Myomatrix recording of motor unit activity in the **mouse bulbospongiosus** muscle evoked by optical stimulation of spinal motor neurons, producing motor unit spikes at latencies between 10-15 msec, consistent with results obtained from traditional fine-wire electrodes in mice ^3^. **(f-j)** Recordings from the **cat soleus (f)** during sensory nerve stimulation, **songbird vocal (ventral syringeal) muscle (g)** and **expiratory muscle (h)** during quiet respiration, **hawkmoth larva dorsal internal medial (DIM) muscle (i)** during fictive locomotion, and **bull frog semimembranosus (SM) muscle (j)** in response to cutaneous (foot) stimulation. Spike times from individual motor units are indicated by colored tick marks under each voltage trace in f-j. Recordings shown in panels (a, c, g, h, i, and j) were collected using bipolar amplification, data in panels (b, d, e, and f) were collected using unipolar recording. See Methods for details of each experimental preparation.

### Myomatrix arrays record from a wide range of muscle groups, species, and behaviors

Myomatrix arrays provide high-resolution EMG recordings across muscle targets and experimental preparations (**Fig. 2**). Beyond the locomotor and postural signals shown in **Figure 1**, Myomatrix recordings from mouse triceps also provided single-unit EMG data during a head-fixed reaching task (**Fig. 2a**). In addition to recording single motor units during these voluntary behaviors, Myomatrix arrays also allow high-resolution recordings from other muscle groups during reflex-evoked muscle activity. **Figure 2b** shows single motor unit EMG data recorded from the superficial masseter (jaw) muscle when reflexive muscle contraction was obtained by passively displacing the jaw of an anesthetized mouse. To extend these methods across species, we collected Myomatrix recordings from muscles of the rat forelimb, obtaining isolated motor units from the triceps during locomotion (**Fig. 2c**) and a digit-flexing muscle in the lower forearm during head-free reaching (**Fig. 2d**). Myomatrix arrays can furthermore isolate motor unit waveforms evoked by direct optogenetic stimulation of spinal motor neurons. **Figure 2e** shows recordings of light evoked spikes in the mouse bulbospongiosus muscle (a pelvic muscle that wraps around the base of the penis in male mice), demonstrating that optogenetic stimulation of the spinal cord evokes spiking in single motor units with millisecond-scale timing jitter (**Fig. 2e**, center) and with latencies (**Fig. 2e**, right) consistent with the latencies of recordings obtained with fine-wire electrodes ^3^. Beyond rodents, simple modifications of the basic electrode array design (**Supplemental Fig. 1f**) allowed us to obtain high-resolution recordings from hindlimb muscles in cats (**Fig. 2f**), vocal and respiratory muscles in songbirds (**Fig. 2g,h**, see also ^27^), body wall muscles in moth larvae (**Fig. 2i**), and leg muscles in frogs (**Fig. 2j**).

In addition to isolating individual motor units, Myomatrix arrays provide stable multi-unit recordings of comparable or superior quality to conventional fine wire EMG. Although single-unit recordings are essential to identify individual motor neurons’ contributions to muscle activity ^28,29^, for other lines of inquiry a multi-unit signal is preferred as it reflects the combined activity of many motor units within a single muscle. Although individual Myomatrix channels are often dominated by spike waveforms from one or a small number of motor units (**Fig 1b**), other channels reflect the combined activity of multiple motor units as typically observed in fine-wire EMG recordings ^30^. As shown in **Supplemental Figure 3a and b**, these multi-unit Myomatrix signals are stable over multiple weeks of recordings, similar to the maximum recording longevity reported for wire-based systems in mice and exceeding the 1-2 weeks of recording more typically obtained with wire electrodes in mice ^1,24,26^, and with significantly greater recording quality than that obtained from wire electrodes at comparable post-implantation timepoints (**Supplemental Fig. 3d**).

To record from larger muscles than those described above, we also created designs targeting the forelimb and shoulder muscles of rhesus macaques (**Fig. 3**). Although fine wire electrodes have been used to isolate individual motor units in both humans and monkeys ^14,31^, and skin-surface electrode arrays robustly record motor unit populations in human subjects ^15,32^, this resolution is limited to isometric tasks – that is, muscle contraction without movement – due to the sensitivity of both fine-wire and surface array electrodes to electrical artifacts caused by body movement. For ease of insertion into larger muscles, we modified the “thread” design used in our mouse arrays so that each Myomatrix array could be loaded into a standard hypodermic syringe and injected into the muscle (**Supplemental Fig. 1g,i**), inspired by earlier work highlighting the performance of injectable arrays in primates ^14,33,34^. As shown in **Figure 3a-d**, injectable Myomatrix arrays yielded motor unit recordings during arm movements. Tick marks in **Figure 3d** show the activity of 13 motor units recorded simultaneously during a single trial in which a monkey was cued (by a force perturbation which causes an extension of the elbow joint) to reach to a target. In contrast to the ensemble of spike times obtained with a Myomatrix probe, conventional fine-wire EMGs inserted into the same muscle (in a separate recording session) yielded only a single trace reflecting the activity of an unknown number of motor units (**Fig 3e**, middle). Moreover, although fine-wire EMG signals varied across trials (**Fig. 3e**, bottom), the lack of motor unit resolution makes it impossible to assess how individual motor units vary (and co-vary) across trials. In contrast, Myomatrix recordings provide spiking resolution of multiple individual motor units across many trials, as illustrated in **Figure 3f**.

**Figure 3:**
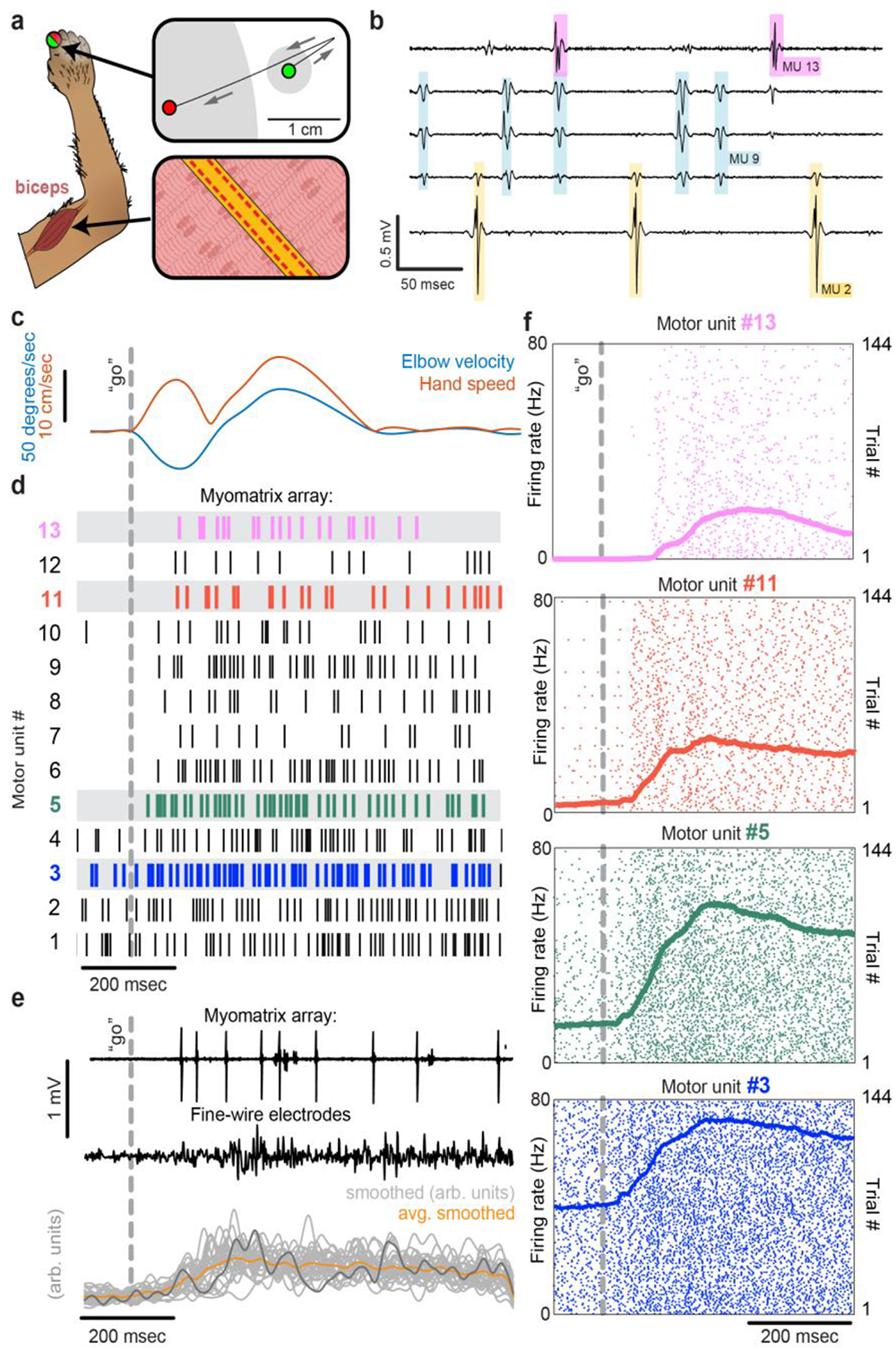
Motor unit recordings during active movement in primates. **(a)** An injectable version of the Myomatrix array (**Supplemental Fig. 1g**) was inserted percutaneously (**Supplemental Fig. 1i**) into the right biceps of a rhesus macaque performing a cued reaching task. Green and red dots: reach start and endpoints, respectively; grey regions: start and target zones. **(b)** Recording from five of 32 unipolar channels showing spikes from three individual motor units isolated from the multichannel recording using Kilosort (**Supplemental Fig. 2**). **(c)** At trial onset (dotted line), a sudden force perturbation extends the elbow, signaling the animal to reach to the target. **(d)** Spike times (tick marks) from 13 simultaneously recorded motor units. **(e)** Example voltage data from a Myomatrix array (top) and traditional fine-wire EMG (middle, bottom) collected from the same biceps muscle in the same animal performing the same task, but in a separate recording session. Gray traces (bottom) show smoothed EMG data from the fine-wire electrodes in all trials, orange trace shows trial-averaged smoothed fine-wire EMG, dark gray trace represents the fine-wire trial shown at middle. **(f)** Spike times of four motor units (of the 13 shown in **d**) recorded simultaneously over 144 trials.

### Myomatrix arrays record stably over time and with minimal movement artifact

Myomatrix arrays provide EMG recordings that are resistant to movement artifacts and stable over time. In recordings from rodents during locomotion (**Fig. 1b**, **Fig. 2c**), we did not observe voltage artifacts at any point in the stride cycle (e.g. when the paw of the recorded limb first touches the ground in each stride cycle; **Fig. 2c**, black arrowhead). Movement artifacts were similarly absent during active arm movements in monkeys (peak fingertip speeds ∼15 cm/sec; peak elbow angle velocity ∼40 deg/sec, **Fig 3b,e**), during passive jaw displacement in anesthetized mice (**Fig. 2b**), or in the other anesthetized preparations shown in **Figure 2**. Individual motor units could typically be isolated for the entire duration of an acute or chronic recording session. In triceps recordings during locomotion in rodents (**Fig. 1**, **Fig. 2c**), isolated motor units were recorded for up to 4,791 stride cycles (mice) or 491 stride cycles (rats) during continuous recording sessions lasting 10-60 minutes. Myomatrix recordings in behaving nonhuman primates were similarly long-lived, as in the dataset shown in **Figure 3**, where single-unit isolation was maintained across 1,292 reaching trials collected over 97 minutes. In each of these datasets, the duration of single-unit EMG recording was limited by the willingness of the animal to continue performing the behavior, rather than a loss of signal isolation. Recordings in acute preparations were similarly stable. For example, the songbird dataset shown in **Figure 2g** includes single-unit data from 8,101 respiratory cycles collected over 74 minutes, and, like the other acute recordings shown in **Figure 2**, recordings were ended by the experimenter rather than because of a loss of signal from individual motor units.

The diversity of applications presented here demonstrates that Myomatrix arrays can obtain high-resolution EMG recordings across muscle groups, species, and experimental conditions including spontaneous behavior, reflexive movements, and stimulation-evoked muscle contractions. Although this resolution has previously been achieved in moving subjects by directly recording from motor neuron cell bodies in vertebrates ^35–37^ and by using fine-wire electrodes in moving insects ^38,39^, both methods are extremely challenging and can only target a small subset of species and motor unit populations. Exploring additional muscle groups and model systems with Myomatrix arrays will allow new lines of investigation into how the nervous system executes skilled behaviors and coordinates the populations of motor units both within and across individual muscles. These approaches will be particularly valuable in muscles in which each motor neuron controls a very small number of muscle fibers, allowing fine control of oculomotor muscles in mammals as well as vocal muscles in songbirds (**Fig. 2g**), in which most individual motor neurons innervate only 1-3 muscle fibers ^40^. Of further interest will be combining high-resolution EMG with precise measurement of muscle length and force output to untangle the complex relationship between neural control, body kinematics, and muscle force that characterizes dynamic motor behavior. Similarly, combining Myomatrix recordings with high-density brain recordings or targeted manipulations of neural activity can reveal how central circuits shape and reshape motor activity and – in contrast to the multi-unit signals typically obtained from traditional EMG in animals – reveal how neural dynamics in cortical, subcortical, and spinal circuits shape the spiking patterns of individual motor neurons.

Applying Myomatrix technology to human motor unit recordings, particularly by using the minimally invasive injectable designs shown in **Figure 3** and **Supplemental Figure 1g,i**, will create novel opportunities to diagnose motor pathologies and quantify the effects of therapeutic interventions in restoring motor function. Moreover, because Myomatrix arrays are far more flexible than the rigid needles commonly used to record clinical EMG, our technology might significantly reduce the risk and discomfort of such procedures while also greatly increasing the accuracy with which human motor function can be quantified. This expansion of access to high-resolution EMG signals – across muscle groups, species, and behaviors – is the chief impact of the Myomatrix project.

## Methods

### Myomatrix array fabrication

The microfabrication process (schematized in **Supplemental Fig. 1a**) consists of depositing and patterning a series of polymer (polyimide) and metal (gold) layers, using a combination of spin coating, photolithography, etching, and evaporation processes, as described previously ^27,41,42^. These methods allow very fine pitch escape routing (<10 µm spacing between the thin “escape” traces connecting electrode contacts to the connector), spatial alignment between the multiple layers of polyimide and gold that constitute each device, and precise definition of “via” pathways that connect different layers of the device. Once all the metal and polyimide layers have been deposited and patterned on carrier wafers, the gold EMG recording electrode sites are formed by removing the top polyimide layer over each electrode site using reactive ion etching process (O2 and SF6 plasma, 5:1 ratio). Electrode sites are then coated with a conductive polymer, PEDOT:PSS (Poly(3,4-ethylenedioxythiophene)-poly(styrene-sulfonate)), ^43–45^ to reduce the electrode impedance ^46^. PEDOT:PSS was deposited on the electrode contacts to a thickness of 100 nm using spin coating, resulting in final electrode impedances of 5 kOhms or less (100 x 200 um electrode sites). Once all layers have been deposited on the carrier wafer, the wafer is transferred to an Optec Femtosecond laser system, which is used to cut the electrode arrays into the shape/pattern needed based on the target muscle group and animal species. The final device thickness was ∼40 µm for the injectable (primate forelimb) design and ∼20 µm for all other design variants. The final fabrication step is bonding a high-density connector (Omnetics, Inc.) to the surface of the electrode array using a Lambda flip-chip bonder (Finetech, Inc.). This fabrication pipeline allows the rapid development and refinement of multiple array designs (**Supplemental Fig. 1c-g**).

### Myomatrix array implantation

For chronic EMG recording in mice and rats (**Fig. 1**, **Fig. 2a, c, d**), arrays such as those shown in **Supplemental Figure 1c-f** were implanted by first making a midline incision (approximately 10 mm length) in the scalp of an anesthetized animal. The array’s connector was then secured to the skull using dental cement (in some cases along with a headplate for later head-fixed chronic recordings), and the electrode array threads were routed subcutaneously to a location near the target muscle or muscles (**Supplemental Fig. 1h**). In some electrode array designs, subcutaneous routing is facilitated with “pull-through tabs” that can be grasped with a forceps to pull multiple threads into position simultaneously. For some anatomical targets a small additional incision was made to allow surgical access to individual muscles (e.g. a 2-5 mm incision near the elbow to facilitate implantation into the biceps and/or triceps muscles). Once each thread has been routed subcutaneously and positioned near its target, any pull-through tabs are cut off with surgical scissors and discarded. Each thread can then either be sutured to the surface of the thin sheet of elastic tissue that surrounds muscles (“epimysial attachment”) or inserted into the muscle using a suture needle (“intramuscular implantation”). For epimysial attachment, each electrode thread is simply sutured to the surface of each muscle (suture sizes ranging from 6-0 to 11-0) in one of the proximal suture holes (located on the depth-restrictor tabs) and one of the distal suture holes. For intramuscular implantation (**Supplemental Fig. 1h**), a suture (size 6-0 to 11-0 depending on anatomical target) is tied to the distal-most suture hole. The needle is then passed through the target muscle and used to pull the attached array thread into the muscle. In some designs, a “depth-restrictor tab” (**Supplemental Fig. 1d**) prevents the thread from being pulled any further into the muscle, thereby limiting the depth at which the electrodes are positioned within the target muscle. The array is then secured within the muscle by the passive action of the flexible polyimide “barbs” lining each thread and/or by adding additional sutures to the proximal and distal suture holes.

Acute recording in small animals (including rodents, songbirds, cats, frogs, and caterpillars; **Fig. 2b,e-j**) used the same arrays as chronic recordings. However, for both epimysial and intramuscular acute recordings, the Myomatrix array traces were simply placed on or within the target muscle after the muscle was exposed via an incision in the overlying skin of the anesthetized animal (rather than routed subcutaneously from the skull as in chronic applications).

For acute recordings in nonhuman primates, prior to recording, the “tail” of the injectable array (**Supplemental Fig. 1g**) was loaded into a sterile 23-gauge cannula (1” long) until fully seated. The upper half of the cannula bevel, where contact is made with the electrode, was laser-blunted to prevent breakage of the tail ^34^. During insertion (**Supplemental Fig. 1i**), the tail was bent over the top of the cannula and held tightly, and the electrode was inserted parallel to bicep brachii long head muscle fibers at an angle of ∼45 degrees to the skin. Once the cannula was fully inserted, the tail was released, and the cannula slowly removed. After recording, the electrode and tail were slowly pulled out of the muscle together. Insertion and removal of injectable Myomatrix devices appeared to be comparable or superior to traditional fine-wire EMG electrodes (in which a “hook” is formed by bending back the uninsulated tip of the recording wire) in terms of both ease of injection, ease of removal of both the cannula and the array itself, and animal comfort. Moreover, in over 100 Myomatrix injections performed in rhesus macaques, there were zero cases in which Myomatrix arrays broke such that electrode material was left behind in the recorded muscle, representing a substantial improvement over traditional fine-wire approaches, in which breakage of the bent wire tip regularly occurs.^14^.

For all Myomatrix array designs, a digitizing, multiplexing headstage (Intan, Inc.) was plugged into the connector, which was cemented onto the skull for chronic applications and attached to data collection devices via a flexible tether, allowing EMG signals to be collected during behavior. By switching out different headstages, data from the same 32 electrode channels on each Myomatrix array could be recorded either as 32 unipolar channels or as 16 bipolar channels, where each bipolar signal is computed by subtracting the signals from physically adjacent electrode contacts.

### Data analysis: spike sorting

Motor unit action potential waveforms from individual motor units were identified with analysis methods previously used to sort spikes from neural data. In all cases, Myomatrix signals (sampling rate 30 or 40 kHz) were first band-passed between 350-7,000 Hz. When the voltage trace from a single Myomatrix channel is dominated by a single high-amplitude action potential waveform (as in **Fig. 1b**), single units can be isolated using principal components analysis (PCA) to detect clusters of similar waveforms, as described previously ^22^. As detailed in **Supplemental Figure 2a-d**, this method provides a simple quantitative measure of motor unit isolation by quantifying the overlap between clusters of spike waveforms in the space of the first two principal components.

In other cases (as in **Fig. 1c**), the spikes of individual motor units appear on multiple channels and/or overlap with each other in time, requiring a more sophisticated spike sorting approach to identifying the firing times of individual motor units. We therefore adapted Kilosort version 2.5 ^2,47^ and wrote custom MATLAB and Python code to sort waveforms into clusters arising from individual motor units (**Supplemental Fig. 2e-h**). Our modifications to Kilosort reflect the different challenges inherent in sorting signals from neurons recorded with Neuropixels probes and motor units recorded with Myomatrix arrays ^14^. These modifications include the following:

> <u>Modification of spatial masking</u>: Individual motor units contain multiple muscle fibers (each of which is typically larger than a neuron’s soma), and motor unit waveforms can often be recorded across spatially distant electrode contacts as the waveforms propagate along muscle fibers. In contrast, Kilosort – optimized for the much more local signals recorded from neurons – uses spatial masking to penalize templates that are spread widely across the electrode array. Our modifications to Kilosort therefore include ensuring that Kilosort search for motor unit templates across all (and only) the electrode channels inserted into a given muscle. In the Github repository linked above, this is accomplished by setting parameter nops.sigmaMask to infinity, which effectively eliminates spatial masking in the analysis of the 32 unipolar channels recorded from the injectable Myomatrix array schematized in **Supplemental Figure 1g**. In cases including chronic recording from mice where only a single 8-contact thread is inserted into each muscle, a similar modification can be achieved with a finite value of nops.sigmaMask by setting parameter NchanNear, which represents the number of nearby EMG channels to be included in each cluster, to equal the number of unipolar or bipolar data channels recorded from each thread. Finally, note that in all cases Kilosort parameter NchanNearUp (which defines the maximum number of channels across which spike templates can appear) must be reset to be equal to or less than the total number of Myomatrix data channels.
>
> <u>Allowing more complex spike waveforms</u>: We also modified Kilosort to account for the greater duration and complexity (relative to neural spikes) of many motor unit waveforms. In the code repository linked above, Kilosort 2.5 was modified to allow longer spike templates (151 samples instead of 61), more spatiotemporal PCs for spikes (12 instead of 6), and more left/right eigenvector pairs for spike template construction (6 pairs instead of 3) to account for the greater complexity and longer duration of motor unit action potentials ^14^ compared to the neural action potentials for which Kilosort was initially created. These modifications were crucial for improving sorting performance in the nonhuman primate dataset shown in **Figure 3**, and in a subset of the rodent datasets (although they were not used in the analysis of mouse data shown in **Fig. 1** and **Supplemental Fig. 2a-f**).

We therefore used Kilosort version 2.5 ^2,47^ and custom MATLAB and Python code to sort waveforms into clusters arising from individual motor units (**Supplemental Fig. 2e-h**). Kilosort 2.5 was modified to allow longer spike templates (151 samples instead of 61), more spatiotemporal PCs for spikes (12 instead of 6), and more left/right eigenvector pairs for spike template construction (6 pairs instead of 3) to account for the greater complexity and longer duration of motor unit action potentials ^14^ compared to the neural action potentials for which Kilosort was initially created.

Individual motor units were identified from “candidate” units by assessing motor unit waveform consistency, SNR, and spike count, by inspecting auto-correlograms to ensure that each identified units displayed an absolute refractory period of less than 1 msec, and by examining cross-correlograms with other sorted units to ensure that each motor unit’s waveforms were being captured by only one candidate unit. Candidate units with inconsistent waveforms or >1% of inter-spike intervals above 1 msec were discarded. Candidate units with highly similar waveform shapes and cross-correlation peaks at lag zero were merged, resulting in sorted units with well-differentiated waveform shapes and firing patterns (**Supplemental Fig. 2e,f**). Our spike sorting code, which includes the above-mentioned modifications to Kilosort, is available at https://github.com/JonathanAMichaels/PixelProcessingPipeline.

Our approach to spike sorting shares the same ultimate goal as prior work using skin-surface electrode arrays to isolate signals from individual motor units but pursues this goal using different hardware and analysis approaches. A number of groups have developed algorithms for reconstructing the spatial location and spike times of active motor units ^18,48^ based on skin-surface recordings, in many cases drawing inspiration from earlier efforts to localize cortical activity using EEG recordings from the scalp ^49^. Our approach differs substantially. In Myomatrix arrays, the close electrode spacing and very close proximity of the contacts to muscle fibers ensure that each Myomatrix channel records from a much smaller volume of tissue than skin-surface arrays. This difference in recording volume in turn creates different challenges for motor unit isolation: compared to skin-surface recordings, Myomatrix recordings include a smaller number of motor units represented on each recording channel, with individual motor units appearing on a smaller fraction of the sensors than typical in a skin-surface recording. Because of this sensor-dependent difference in motor unit source mixing, different analysis approaches are required for each type of dataset. Specifically, skin-surface EMG analysis methods typically use source-separation approaches that assume that each sensor receives input from most or all of the individual sources within the muscle as is presumably the case in the data. In contrast, the much sparser recordings from Myomatrix are better decomposed using methods like Kilosort, which are designed to extract waveforms that appear only on a small, spatially restricted subset of recording channels.

### Additional recording methods – mouse forelimb muscle

All procedures described below were approved by the Institutional Animal Care and Use Committee at Emory University (data in **Fig. 1c**, **Supplemental Fig 3**) or were approved by European Committee Council Directive, the Animal Care and Users Committee of the Champalimaud Neuroscience Program, and the Portuguese National Authority for Animal Health (data in **Fig. 1b, Supplemental Fig. 2e**). Individual Myomatrix threads were implanted in the triceps muscle using the “intramuscular” method described above under isoflurane anesthesia (1-4% at flow rate 1 L/min). EMG data were then recorded either during home cage exploration or while animals walked on a custom-built linear treadmill ^50^ at speeds ranging from 15-25 cm/sec. A 45° angled mirror below the treadmill allowed simultaneous side and bottom views of the mouse ^6^ using a single monochrome usb3 camera (Grasshopper3, Teledyne FLIR) to collect images 330 frames per second. We used DeepLabCut ^51^ to track paw, limb, and body positions. These tracked points were used to identify the stride cycles of each limb, defining stance onset as the time at which each paw contacts the ground and swing onset as the time when each paw leaves the ground.

### Additional recording methods – mouse orofacial muscle

All procedures described below were approved by The Institutional Animal Care and Use Committee at Johns Hopkins University. Individual Myomatrix threads were implanted on the masseter muscle using the “epimysial” method described above. A ground pin was placed over the right visual cortex. As described previously ^52^, EMG signals and high-speed video of the orofacial area were recorded simultaneously in head-fixed animals under isoflurane anesthesia (0.9-1.5% at flow rate 1L/min). During data collection, the experimenter used a thin wooden dowel to gently displace the mandible to measure both jaw displacement and muscle activity from the jaw jerk reflex. Jaw kinematics were quantified using a high-speed camera (PhotonFocus DR1-D1312-200-G2-8) at 400 frames per second using an angled mirror to collect side and bottom views simultaneously. Jaw displacement was quantified by tracking eleven keypoints along the jaw using DeepLabCut ^51^.

### Additional recording methods – rat forelimb muscle

All procedures described below were approved by The Institutional Animal Care and Use Committee at Emory University. Anesthesia was induced with an initial dose of 4% isoflurane in oxygen provided in an induction chamber with 2 L/minute rate and maintained with 3% isoflurane at 1 L/minute. Following this, rats received a subcutaneous injection of 1mg/kg Meloxicam, a subcutaneous injection of 1% Lidocaine and topical application of lidocaine ointment (5%) at each incision site. Myomatrix threads were implanted in the triceps muscle using the “intramuscular” method. EMG data were then recorded while animals walked on a treadmill at speeds ranging from 8-25 cm/sec. Kinematics were quantified using a circular arrangement of four high-speed FLIR Black Fly S USB3 cameras (BFS-U3-16S2M-CS, Mono), each running at 125 FPS. We used DeepLabCut to label pixel locations of each of ten anatomical landmarks on the limbs and body, which we then transformed into 3D cartesian coordinates using Anipose ^51,53^. We then defined the onset of each swing/stance cycle by using local minima in the rat’s forelimb endpoint position along the direction of locomotion.

### Additional recording methods – rat forelimb muscle

All procedures described below were approved by The Institutional Animal Care and Use Committee at Johns Hopkins University. Prior to electrode implantation, rodents were trained for 4-6 weeks to perform a single pellet reach task ^54^. Rodents were food restricted for 17-18 hours prior to training. During the task, rats used the right arm to reach for sucrose pellets through a vertical slit (width = 1 cm) in a custom-built acrylic chamber. Individual MyoMatrix threads were implanted on the right flexor digitorum profundus (FDP) muscle using the “epimysial” method described above under 2.0-3.0% isoflurane anesthesia in oxygen gas. The arrays were then connected to data collection hardware via a flexible tether and EMG data were recorded while unrestrained animals performed the reaching task. Kinematics were quantified using a smartphone camera running at 60 fps. Two raters then manually labelled the frames of grasp onset. Grasp initiation was defined when a frame of full digit extension was immediately followed by a frame of digit flexion.

### Additional recording methods – mouse pelvic muscle

All experimental procedures were carried out in accordance with the guidelines of the European Committee Council Directive and were approved by the Animal Care and Users Committee of the Champalimaud Neuroscience Program, the Portuguese National Authority for Animal Health. As described in detail elsewhere ^3^, spinal optogenetic stimulation of the motor neurons innervating the bulbospongiousus muscle (BSM) was performed on 2-3 month old male BL6 mice that had received an injection of a rAAVCAG-ChR2 into the BSM on postnatal day 3-6. Individual Myomatrix threads were implanted in the BSM using the “intramuscular” method described above. During EMG recording an optrode was moved on top of the spinal cord along the rostral-caudal axis while applying optical stimulation pulses (10 msec duration, power 1-15 mW).

### Additional recording methods – songbird vocal and respiratory muscles

All procedures described below were approved by The Institutional Animal Care and Use Committee at Emory University. As described previously ^20,27,42^, adult male Bengalese finches (>90 d old) were anesthetized using intramuscular injections of 40 mg/kg ketamine and 3 mg/kg midazolam injected and anesthesia was maintained using 1-5% isoflurane in oxygen gas. To record from the expiratory (respiratory) muscles, an incision was made dorsal to the leg attachment and rostral to the pubic bone and the electrode array was placed on the muscle surface using the “epimysial” approach described above. To record from syringeal (vocal) muscles, the vocal organ was accessed for electrode implantation via a midline incision into the intraclavicular air sac as described previously ^20^ to provide access to the ventral syringeal (VS) muscle located on the ventral portion of the syrinx near the midline.

### Additional recording methods – cat soleus muscle

All procedures described below were approved by The Institutional Animal Care and Use Committee at Temple University. As described previously ^55^, an adult female cat was provided atropine (0.05 mg/kg IM) and anesthetized with isoflurane (1.5–3.5% in oxygen), during which a series of surgical procedures were performed including L3 laminectomy, implantation of nerve cuffs on the tibial and sural nerve, and isolation of hindlimb muscles. Individual Myomatrix threads were implanted in hindlimb muscles using the “intramuscular” method described above. Following these procedures, a precollicular decerebration was performed and isoflurane was discontinued. Following a recovery period, the activity of hindlimb motor units were recorded in response to electrical stimulation of either the contralateral tibial nerve or the ipsilateral sural nerve.

### Additional recording methods – hawkmoth larva (caterpillar) body wall muscle

EMG recordings were obtained from 5th instar larvae of the tobacco hornworm Manduca sexta using a semi-intact preparation called the “flaterpillar” as described previously (Metallo, White, and Trimmer 2011). Briefly, after chilling animals on ice for 30 minutes, an incision was made along the cuticle, allowing the nerve cord and musculature to be exposed and pinned down in a Sylgard dish under cold saline solution. This preparation yields spontaneous muscle activity (fictive locomotion) which was recorded from the dorsal intermediate medial (DIM) muscle using the “epimysial” method described above, with the modification that sutures were not used to hold the muscle in place.

### Additional recording methods – frog hindlimb muscles

Spinal bullfrogs were prepared under anesthesia in accordance with USDA and PHS guidelines and regulations following approval from the Institutional Animal Care and Use Committee at Drexel University as described previously^56^. Bull frogs were anesthetized with 5% tricaine (MS-222, Sigma), spinalized, and decerebrated. The frog was placed on a support and Myomatrix arrays were implanted into the semimembranosus (SM) hindlimb muscle using the “intramuscular” method described above. Epidermal electrical stimulation at the heel dorsum (500 msec train of 1 msec, 5V biphasic pulses delivered at 40 Hz) or foot pinch was used to evoke reflexive motor activity.

### Additional recording methods – rhesus macaque forelimb muscle

All procedures described below were approved by The Institutional Animal Care and Use Committee at Western University. One male rhesus monkey (Monkey M, Macaca mulatta, 10 kg) was trained to perform a range of reaching tasks while seated in a robotic exoskeleton (NHP KINARM, Kingston, Ontario). As described previously ^57,58^, this robotic device allows movements of the shoulder and elbow joints in the horizontal plane and can independently apply torque at both joints. Visual cues and hand feedback were projected from an LCD monitor onto a semi-silvered mirror in the horizontal plane of the task and direct vision of the arm was blocked with a physical barrier.

An injectable Myomatrix array (**Supplemental Fig. 1g**) was inserted percutaneously as shown in **Supplemental Figure 1i**. Then, using his right arm, Monkey M performed a reaching task similar to previous work ^57^. On each trial the monkey waited in a central target (located under the fingertip when the shoulder and elbow angles were 32° and 72°, respectively; size = 0.6 cm diameter) while countering a constant elbow load (-0.05 Nm). The monkey was presented with one of two peripheral goal targets (30/84° and 34/60° shoulder/elbow, 8cm diameter), and after a variable delay (1.2-2s) received one of two unpredictable elbow perturbations (±0.15Nm step-torque) which served as a go cue to reach to the goal target. At the time of perturbation onset, all visual feedback was frozen until the hand remained in the goal target for 800ms, after which a juice reward was given. On 10% of trials no perturbation was applied, and the monkey had to maintain the hand in the central target. In addition to Myomatrix injectables, we acquired bipolar electromyographic activity from nonhuman primates using intramuscular fine-wire electrodes in the biceps brachii long head as described previously ^59^, recording in this instance from the same biceps muscle in the same animal from which we also collected Myomatrix data, although in a separate recording session. Fine-wire electrodes were spaced ∼8 mm apart and aligned to the muscle fibers, and a reference electrode was inserted subcutaneously in the animal’s back. Muscle activity was recorded at 2,000 Hz, zero-phase bandpass filtered (25–500 Hz, fourth order Butterworth) and full-wave rectified.

## Data and code availability

A data archive including two EMG datasets recorded with Myomatrix arrays from behaving animals is available at doi.org/10.5061/dryad.66t1g1k70. This archive includes the mouse triceps data shown in **Figure1 (b,d)** and **Supplemental Figure 2(a,b,e,f)**, the rhesus macaque biceps data shown in **Figure 3** and **Supplemental Figure 2g**, and associated metadata.

---

**Supplemental Figure 1:**
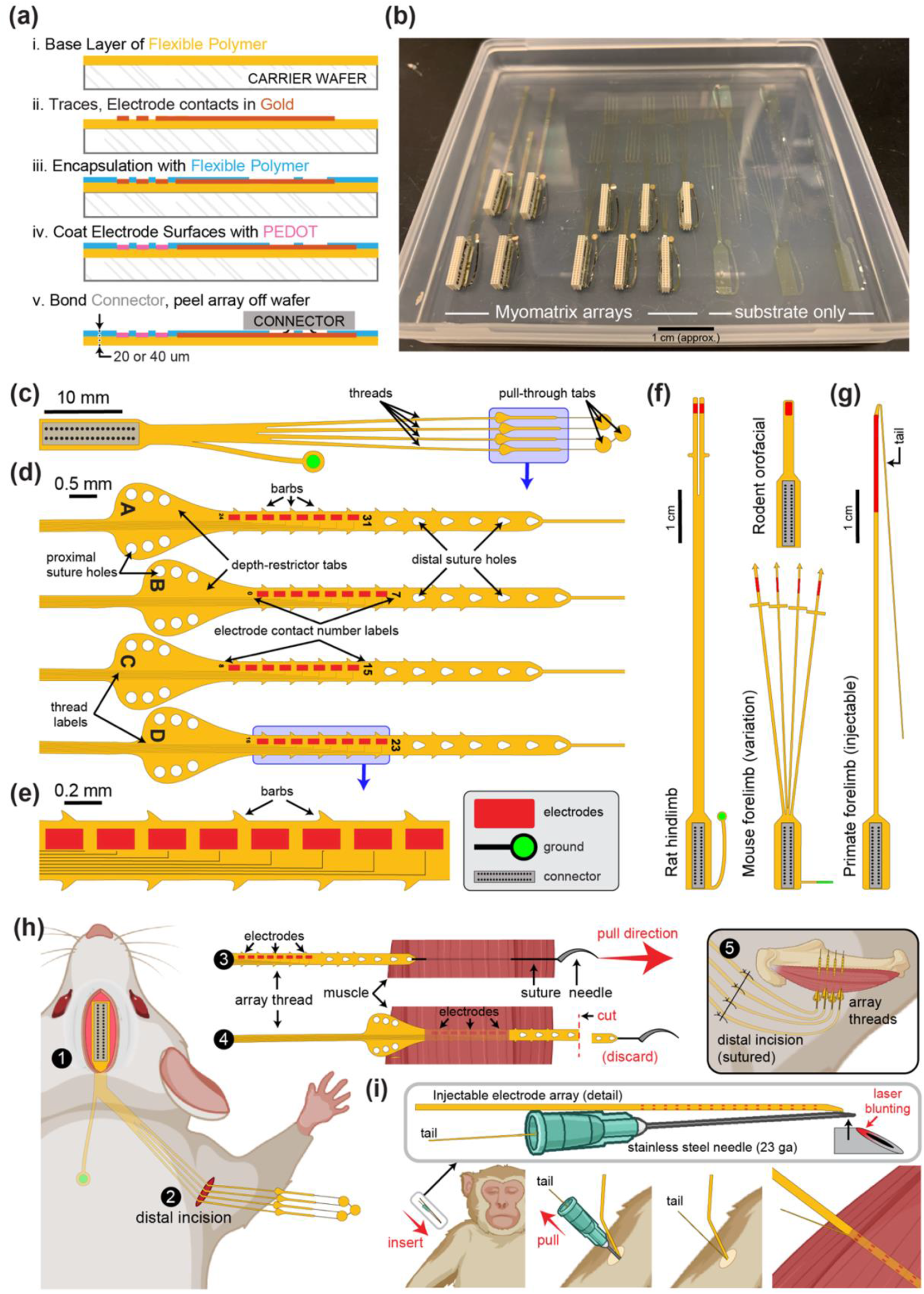
Myomatrix fabrication, design variations, and implantation. **(a)** Workflow for electrode array fabrication. Layers of insulating polymer (polyimide) and conductive metal (gold) are successively deposited on a carrier wafer to form a flexible, 20 or 40 μm thick electrode array of gold electrode contacts, which receive a surface treatment of PEDOT to improve recording properties (see Methods). Electrodes are connected via thin gold traces to a receiving pad for a high-density connector (Omnetics Inc.) which is then bonded to the array. The completed array is then peeled off the carrier wafer. **(b)** Photo showing two different Myomatrix designs (left) as well as “blank” arrays comprised only of the flexible polyimide substrate for surgical practice and design optimization. **(c-e)** Expanded views of the electrode array also shown in Figure 1 of the main manuscript text, which has four “threads” each bearing eight electrode contacts. This array design can be used for either acute or chronic recordings. For chronic implantation, the surgeon grasps the “pull-through tabs” when tunneling the threads subcutaneously. For intramuscular implantation in either acute or chronic settings, a needle is used to pull each thread through the target muscle. In this use case, the “depth-restrictor tabs” prevent the thread from being pulled any further into the muscle, thereby determining the depth of the electrode contacts within the muscle. **(d,e)** Detail views highlighting sub-millimeter features used to increase electrode stability within the muscle (barbs, suture holes) and labels to indicate which channel/thread labels have been implanted in which muscles. **(f,g)** Design variations. The fabrication process shown in **(a)** can easily be modified to alter size and shape of the electrode array. Each Myomatrix design in **f** and **g** has 32 electrode contacts. **(f)** Two array designs customized for chronic recording applications in different muscle groups in rodents. **(g)** Injectable array for recording forelimb muscles in nonhuman primates. **(h)** For chronic implantation in mice, the connector end of the array is attached to the skull using dental acrylic (1) and the flexible array threads are then routed subcutaneously to a small distal incision located near the targeted muscle or muscles (2). For intramuscular implantation, the surgeon secures each thread to a suture and needle, which are then inserted through the target area of muscle tissue (3). The surgeon then pulls the suture further through the muscle, eventually drawing the array thread into the muscles such that the depth-restrictor tabs prevent further insertion and ensure that the electrode contacts are positioned at the correct depth within the muscle (4). In contrast, for epimysial (as opposed to intramuscular) implantation, the array threads are sutured to the surface of the muscle fascia rather than being inserted with a suture needle. After all threads are secured to the muscle, the distal incision site is sutured closed (5). **(i)** For percutaneous insertion of injectable arrays, the array’s thin “tail” is loaded into a modified hypodermic syringe ^14,34^. During insertion, the tail is secured by bending it back over the plastic needle holder and securing it with either the surgeon’s fingers or an additional syringe inserted into the cannula. After electrode array insertion the needle is gently pulled out of the muscle, leaving the electrode-bearing part of the array thread within the target muscle for the duration of the recording session. After recording, the electrode and tail are gently pulled out of the muscle together as with injectable fine-wire EMG ^14^.

**Supplemental Figure 2:**
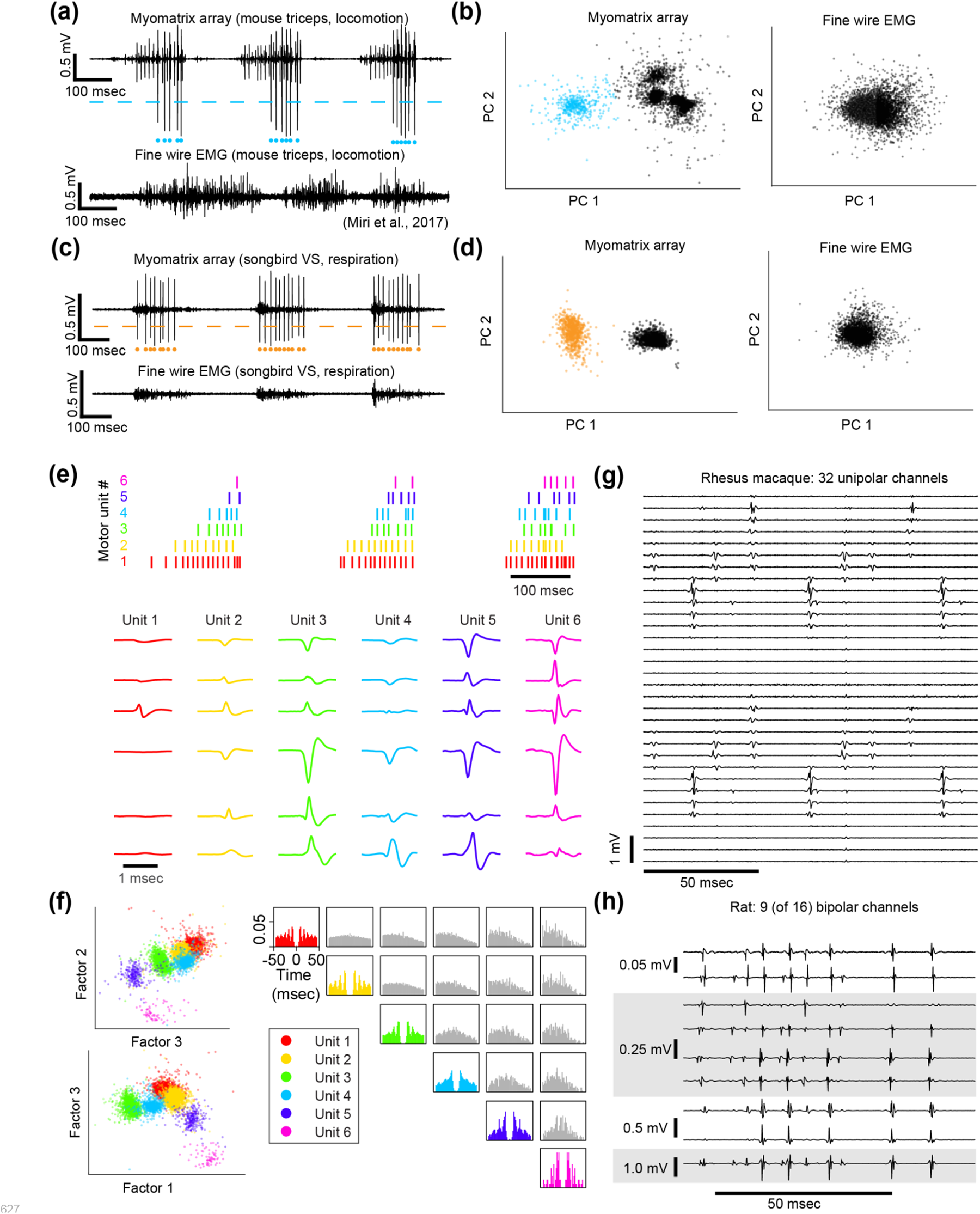
Spike sorting Action potential (“spike”) waveforms from individual motor units can be identified (“sorted”) using analysis methods that are commonly used to sort spikes from neural data. **(a-d)** Single channel spike sorting. In some cases, a single motor unit’s spike will dominate the recording on an individual Myomatrix channel, as shown in an example bipolar recording from mouse triceps during locomotion (**a**, top). In such cases, a simple voltage threshold (dashed line) can be used to isolate spike times of the largest recorded unit (blue dots) from a single channel. In contrast (**a,** bottom), fine-wire EMG typically does not yield isolated single units during active behaviors. **(b)** Single-channel spike sorting using principal components analysis (PCA) of the data shown in (a). Each data point in (b) represents a single voltage waveform represented in the dimensions defined by the first two principal components (PC1 and PC2) of the set of all spike waveforms. As described previously ^22^, k-means clustering can discriminate the waveforms from individual motor units (cyan dots in **a** and **b**) and waveforms from other motor units and/or background noise (black dots in **b**). If one of the clusters has less than 1% overlap with any other cluster (based on fitting each cluster with a 2D Gaussian as described previously) and displays an absolute refractory period (less than 1% of inter-spike intervals less than 1 msec), it is classified as a single unit ^22^. When applied to the Myomatrix data in (a), PCA-based sorting method produced identical spike times as the thresholding method (cyan dots in **a**). In contrast, the same analysis applied to the fine-wire data shown in **a** did not produce any well-isolated clusters in PCA space (**b**, right), indicating that this method could not extract any single motor units. Myomatrix and fine-wire data shown in (a,b) are from the same datasets as the examples shown in main text Figure 1a,b. **(c,d)** Single-channel spike sorting applied to bipolar Myomatrix recordings from the ventral syringeal (VS) muscle, a songbird vocal muscle ^20^. Here again, PCA-based sorting of Myomatrix data method produced identical spike times as the thresholding method (orange dots in **c** and **d**). In contrast, the same analysis applied to fine-wire data recorded from VS shown in **c** did not produce any well-isolated clusters in PCA space (**d**, right), Other plotting conventions for (c,d) are the same as for the mouse data in (a,b). **(e-h)** Multichannel spike sorting using Kilosort. We used Kilosort version 2.5 ^2,47^ and custom MATLAB and Python code to sort waveforms into clusters arising from individual motor units. **(e)** Spike times (top) and mean waveforms (bottom) of six motor units recorded simultaneously from mouse triceps during locomotion (same dataset as Fig. 1 in the main text). Mean waveforms for the six motor units (columns at bottom) are shown from six different EMG channels (rows) and illustrate the distinct pattern of spike waveforms across channel associated with the discharge of each identified motor unit. **(f)** Left, feature space projection of individual waveforms (colored dots), projected onto the space of singular values (“factors”) that describe the space of all recorded waveforms. The clustering of waveforms from kilosort-identified units (colors) further illustrates the distinctness of voltage waveforms assigned to each of the identified motor units. Right, autocorrelograms (colors) and cross correlograms (gray) of the six motor units shown in (e). In addition to examining the consistency of each candidate motor unit’s spike waveforms we also inspected autocorrelations to ensure that each identified unit showed an absolute refractory period (zero or near-zero autocorrelations at lag zero) and that cross-correlograms did not have strong peaks at zero lag (which might indicate the same motor unit being detected by multiple Kilosort clusters). **(g,h)** Myomatrix recordings from nonhuman primate and rat (unipolar and bipolar recordings respectively, same datasets as in main text Fig. 3 **and** Fig. 2c), respectively. These examples (along with the mouse data in main text Fig. 1c) highlight the finding that Myomatrix arrays typically record the same motor unit on multiple channels simultaneously. This redundancy is critical for Kilosort and related methods to isolate single motor unit waveforms, particularly when waveforms from multiple units overlap in time.

**Supplemental Fig 3:**
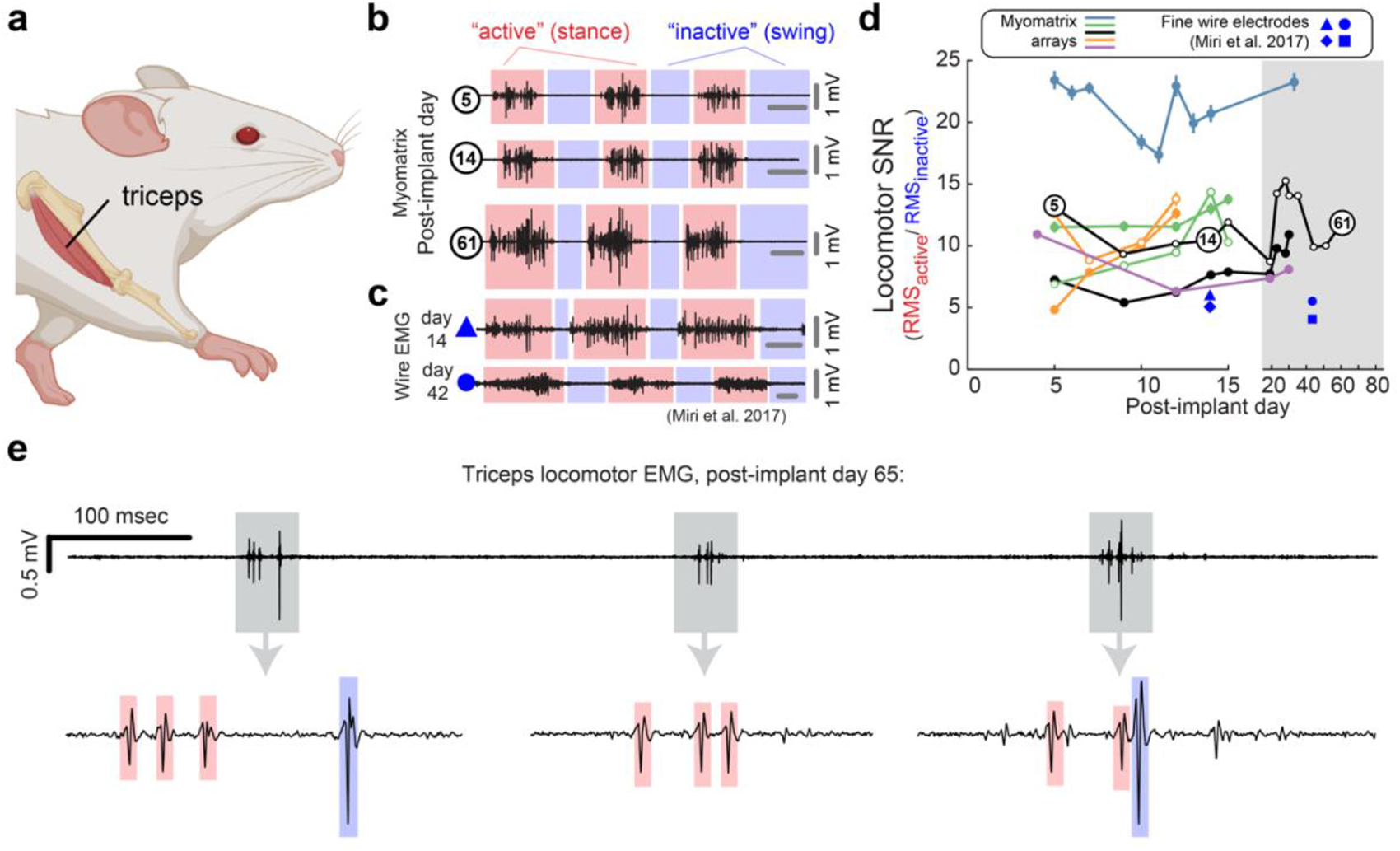
Longevity of Myomatrix recordings In addition to isolating individual motor units, Myomatrix arrays also provide stable multi-unit recordings of comparable or superior quality to conventional fine wire EMG. **(a,b)** Bipolar Myomatrix recordings from the triceps muscle of a mouse recorded during treadmill locomotion over a period 61 days. Colored regions in (b) highlight the “stance” phase (when the paw from the recorded forelimb is in contact with the treadmill surface) and “swing” phase (when the paw is lifted off the treadmill surface). To quantify changes in recording quality over time, we computed a “signal-to-noise ratio (SNR)” for each of each stride cycle as described previously ^21^. Here, the “locomotor SNR” for each swing-stance-cycle is defined as the root mean square (RMS) amplitude of the multi-unit EMG signal during each single stance cycle divided by the RMS of the EMG signal during the immediately subsequent swing phase. **(c)** Fine-wire EMG data recorded from the triceps muscle during locomotion (reproduced with permission from ^1^. Note that all horizontal gray bars in (b,c) represent 100 msec. **(d)** Mean +/-standard error of locomotor SNR across five mouse subjects implanted with Myomatrix arrays. Filled symbols indicate EMG implantation in the right triceps muscle, unfilled symbols indicate EMG implantation in the left triceps. The black trace with unfilled symbols represents the animal whose data are also shown in panel (b). In some cases, error bars are hidden behind plotting symbols. Blue symbols indicate the locomotor SNR from the fine-wire data from ^1^, with each symbol representing a single day’s recording from one of four individual mice. SNR values from Myomatrix arrays are significantly greater than those from fine-wire EMG, both when all data shown in (d) are pooled and when only data from day 14 are included (2-sample KS-test, p=0.002 and p=0.038, respectively). (e) Although individual motor units were most frequently recorded in the first two weeks of chronic recordings (see main text), Myomatrix arrays also isolate individual motor units after much longer periods of chronic implantation, as shown here where spikes from two individual motor units (colored boxes in bottom trace) were isolated during locomotion 65 days after implantation. This bipolar recording was collected from the subject plotted with unfilled black symbols in panel (d).

## Acknowledgements

The participants of the Emory-SKAN Remote Workshop for Advanced EMG Methods, which brought together over 100 researchers from around the world, for their critical feedback on how to improve and refine the electrode technology described here.

Dr. Andrew Miri for helpful discussions and for sharing locomotor EMG data from Miri et al. (2017). Dr. Cinzia Metallo for the initial *Manduca* studies using flexible electrode arrays.

Drs. Gabriela J. Martins and Mariana Correia for project and colony coordination. Dr. Ana Gonçalves for technical assistance in mouse locomotion experiments.

Mattia Rigotti, Margo Shen, Nevin Aresh, and Manikandan Venkatesh for assistance in collecting rat forelimb data. Components of all figures were created using BioRender.com.

## Funding sources

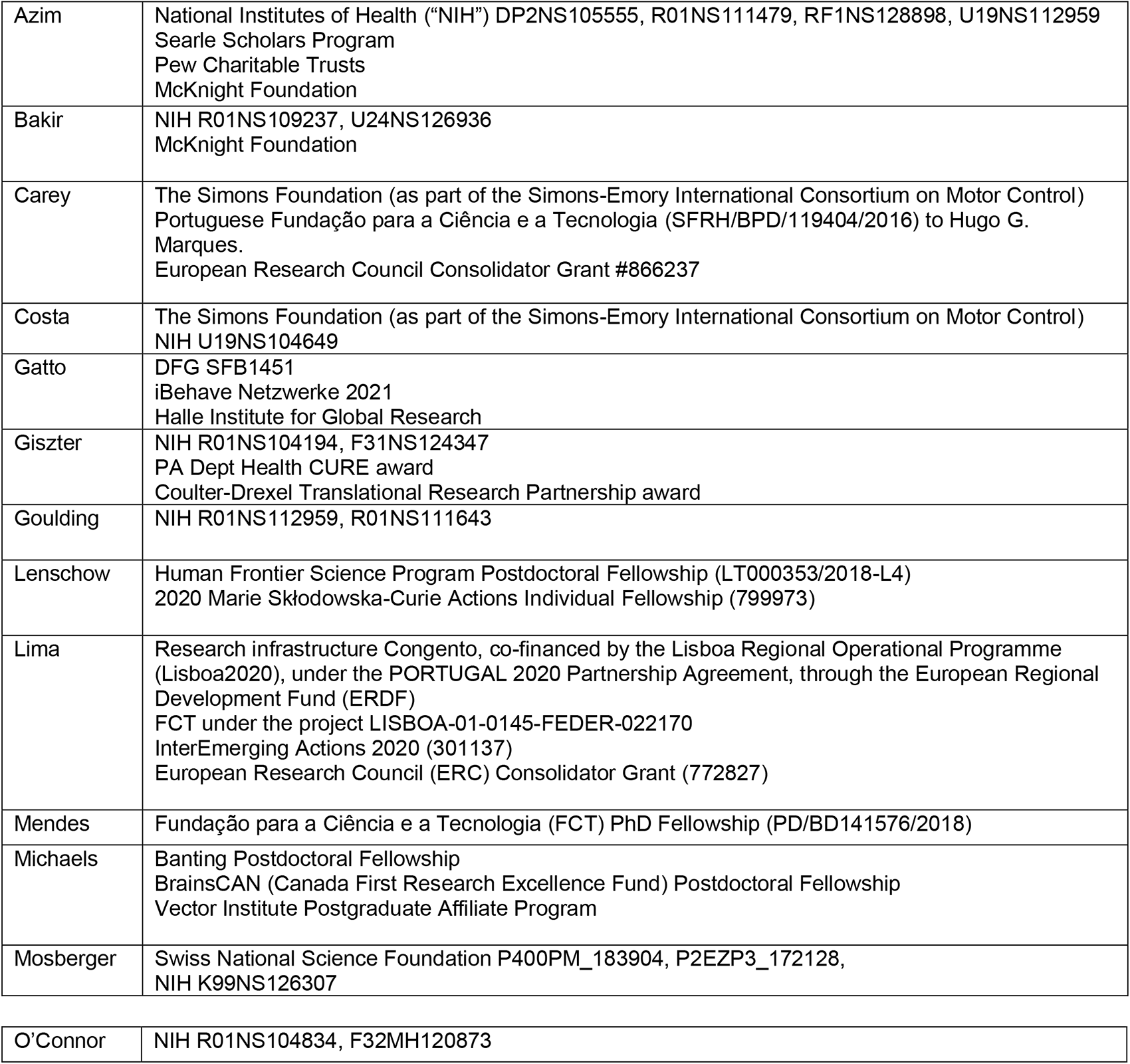
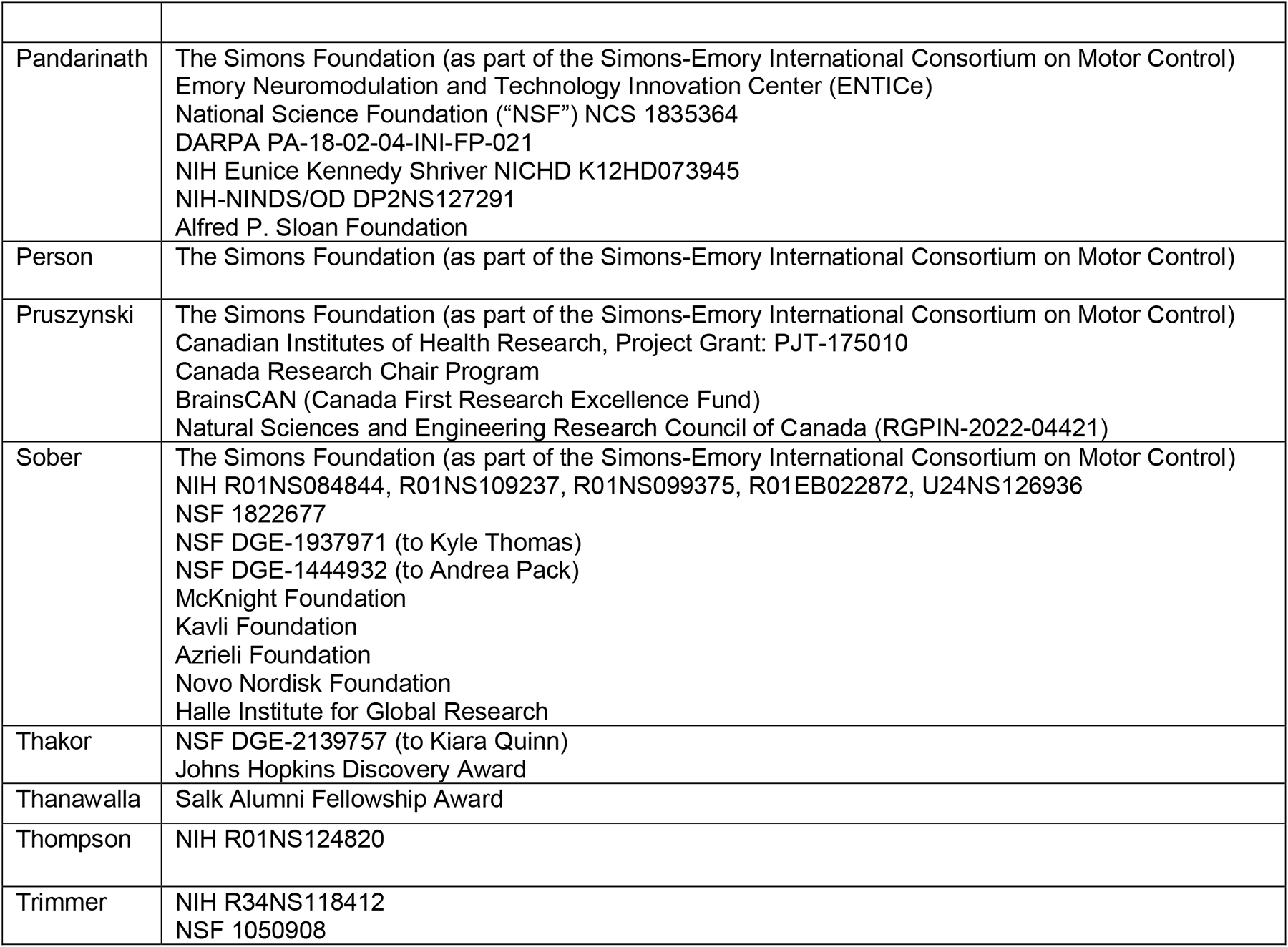

